# One placement, many fields: robust voxel-wise mapping with multichannel TMS

**DOI:** 10.64898/2026.10.24.761539

**Authors:** Ole Numssen, Renxiang Qiu, Victor Souza, Mikael Laine, Renan H. Matsuda, Lena Buchberger, Carla W. Martin, Nico Scherf, Konstantin Weise, Thomas R. Knösche

**Affiliations:** Methods and Development Group Brain Networks, Max Planck Institute for Human Cognitive and Brain Sciences, Leipzig, Germany; Methods and Development Group Neural Data Science and Statistical Computing, Max Planck Institute for Human Cognitive and Brain Sciences, Leipzig, Germany; Center for Scalable Data Analytics and Artificial Intelligence (Scads.AI), Leipzig University, Germany; Department of Neuroscience and Biomedical Engineering, Aalto University, Espoo, Finland; Cortisys Ltd, Espoo, Finland; Edmond and Lily Safra International Institute of Neuroscience, Santos Dumont Institute, Macaíba, Brazil; Leipzig University of Applied Sciences, HTWK, Leipzig, Germany; Institute of Biomedical Engineering and Informatics, Technische Universität Ilmenau, Ilmenau, Germany

## Abstract

Understanding the human brain at the individual level is central to understanding its function and a prerequisite for personalized therapy. Transcranial magnetic stimulation (TMS) can map structure–function relationships at high resolution by regressing trial-wise variation in the TMS-induced electric field (E-field) against a response such as motor evoked potentials (MEPs). This requires distinct E-fields across pulses at different cortical locations, which single-coil TMS (sTMS) achieves by physically repositioning the coil—increasingly reliant on robotic guidance and capped by the coil geometry. Multichannel TMS (mTMS) instead steers the E-field electronically via independent currents in several coils, opening a design space of high variability without physical repositioning.

We adapt determinant-based sequence optimization from discrete sTMS coil-placement sets to the continuous mTMS current space (D-optimal design), identifying pulse sequences whose induced cortical E-fields are as mutually dissimilar as biophysical constraints allow. We benchmarked optimized mTMS sequences against random baselines and optimized sTMS placements in virtual motor mapping across multiple head models, assessing three mTMS array geometries (5-channel planar, 6- and 12-channel spherical), two figure-of-eight sTMS coils, and varying pulse counts, response noise, and cortical depth.

D-optimal mTMS currents substantially outperformed random currents and optimized sTMS at matched pulse counts. The gain was modest without noise but increased significantly with it: at the highest noise level, optimized sTMS failed to converge after 100 pulses (∼20–22 mm error), whereas all mTMS arrays reached median errors at or below 6 mm. mTMS also recovered deep targets better than sTMS (up to 35 mm skin–cortex distance, versus 25 mm) and performed uniformly across the region of interest. In addition, we found that a coil-space scheme based only on the magnetic vector potential, precomputable once per array without per-subject MRI, captured most of the benefit, matching subject-specific optimization at 100 pulses. mTMS thus makes voxel-wise mapping robust throughout a region of interest without coil repositioning.

## 1 Introduction

Transcranial magnetic stimulation (TMS) has become a cornerstone of non-invasive brain stimulation (NIBS) for research, clinical, and therapeutic applications (Pascual-Leone et al., 2000; Valero-Cabré et al., 2017; Di Lazzaro et al., 2021). By delivering brief but strong electromagnetic pulses via a coil positioned on the scalp, TMS can induce action potentials in underlying cortical neurons and thereby perturb local circuits as well as distributed networks (Siebner et al., 2009). This above-threshold perturbation enables causal inferences on brain function by linking controlled cortical stimulation to physiological or behavioral outcomes (Bergmann & Hartwigsen, 2021). To assess the engagement of different cortical areas to a specific cognitive or motor function, the TMS coil has traditionally been repositioned between pulses or stimulation blocks, shifting the peak location of the induced electric field (E-field) across the cortical surface (Thielscher et al., 2015; Rossini et al., 2015; Groppa et al., 2012).

Multichannel or multi-locus TMS (mTMS) extends conventional single-coil TMS (sTMS) by driving multiple coils with independently controlled currents (Ruohonen & Ilmoniemi, 1998), enabling fast electronic steering of intracranial E-fields and a rich space of stimulation patterns without mechanical repositioning (Navarro de Lara et al., 2021; Nieminen & Sinisalo et al., 2022; Tervo & Nieminen et al., 2022; Souza et al., 2022; Souza et al., 2025; Daneshzand et al., 2025a,b; Koponen et al., 2018; Sinisalo & Rissanen et al., 2024). In principle, this flexibility (Soto et al., 2026) supports more efficient cortical mapping, more selective targeting, and adaptive, closed-loop protocols that tailor the induced fields and their dose (Kim et al., 2026; Numssen & Kuhnke et al., 2024) to individual anatomy and functional organization (Bergmann & Hartwigsen, 2021; Numssen & Kuhnke et al., 2024). In practice, however, we still lack a principled framework for deciding which mTMS current patterns to deliver and how to optimally exploit the high-dimensional control space. We recently proposed a focality-first framework that incorporates individual head and brain morphology to optimize mTMS currents for stimulating a particular target and minimizing off-target stimulation, thereby quantifying attainable focality and controllability for specific coil geometries (Numssen & Martin et al., 2026). While this strategy is well suited when the stimulation target is already known, mapping unknown structure-function relationships calls for a different approach: voxel-wise TMS mapping, which uses sets of distinct TMS pulses to map an entire region of interest at once rather than optimizing single targets sequentially (Bungert et al., 2017; Numssen et al., 2021; Weise & Numssen et al., 2023; Schultheiss & Turi et al., 2025; Vetter et al., 2025). Only recently, the TMS-induced E-field for a specific coil and placement has become accurately modellable for individual head and brain anatomies on standard hardware, substantially advancing our understanding of the cortical effects of TMS (Thielscher et al., 2015; Wartman et al., 2025; Worbs et al., 2024; Cao et al., 2024; Soto et al., 2026; Siebner et al., 2022). Leveraging this, voxel-wise TMS mapping was introduced as a structure-function mapping framework that integrates E-field simulations across multiple coil positions and orientations to localize functional representations with high spatial precision (Weise & Numssen et al., 2020; Numssen et al., 2021; Weise & Numssen et al., 2023). Conceptually analogous to mass-univariate functional MRI or source-space EEG/MEG analyses, voxel-wise TMS mapping statistically associates voxel-wise stimulation strength with outcome measures such as motor evoked potentials (MEPs), turning trial-wise E-field variation into cortical effect maps of TMS (Fig. 1; Weise & Numssen et al., 2020; Laine et al., 2026; Vetter et al., 2025). Recent studies have validated and extended this regression-based approach beyond primary motor cortex, for example, to spatial attention (Jing et al., 2023) and heart-brain coupling (Feng & Martin et al., 2026; Feng et al., 2026), highlighting its potential to shift TMS from a coil-centric to a data-driven, E-field-centric mapping technique (Jing et al., 2023; Kuhnke et al., 2025).

**Figure 1:**
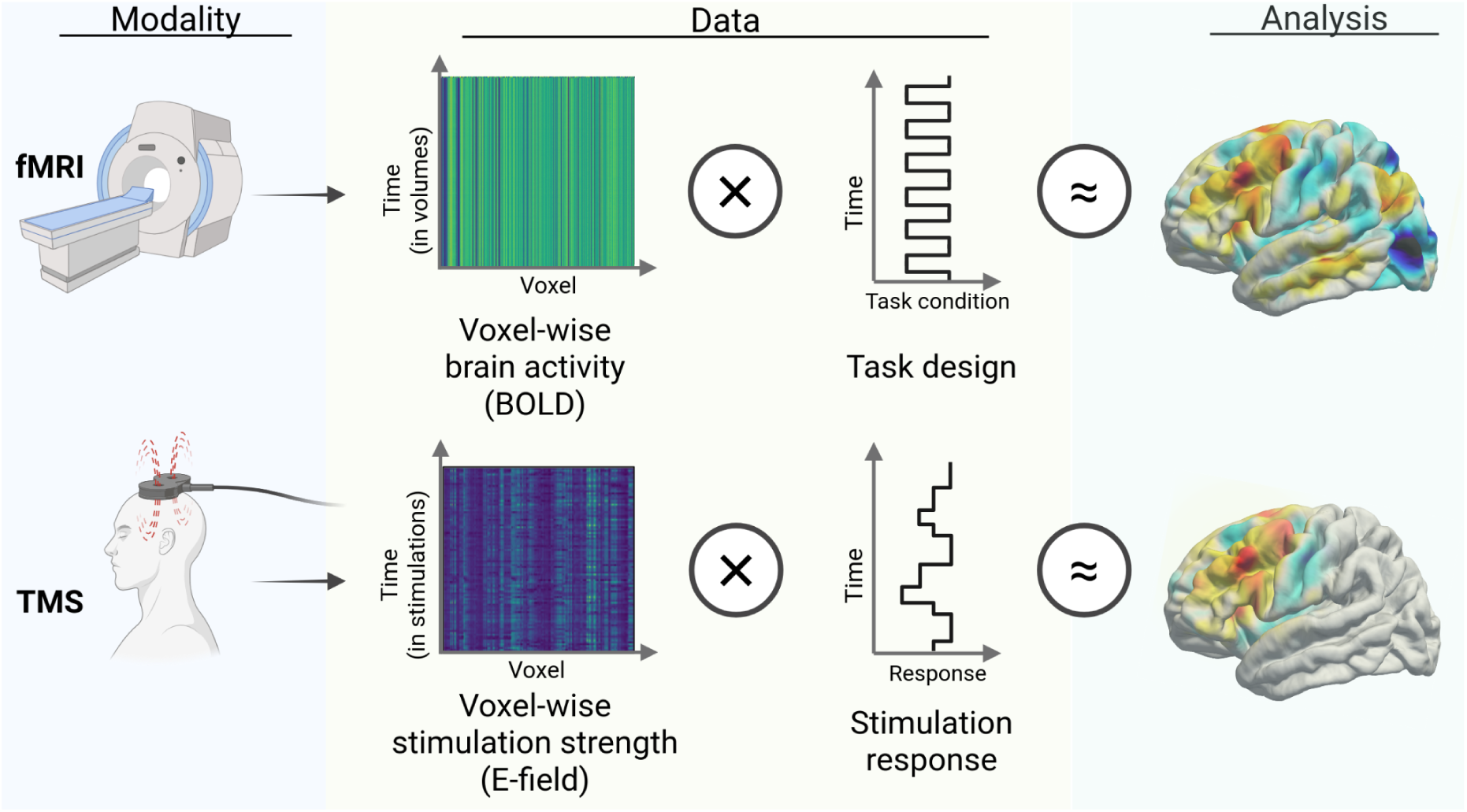
Voxel-wise TMS mapping and its analogy to univariate fMRI analysis. Top: In standard mass-univariate functional MRI analyses, a task design is fit to voxel-wise brain activity over time, i.e., volumes. This analysis aims to answer the question “*Where does brain activity fit best to the task-design?”*. Bottom: In voxel-wise TMS mapping, voxel-wise brain stimulation, i.e., the TMS-induced E-field from different coil placements, is fit to the stimulation response, such as motor evoked potentials (MEPs) or reaction time changes. Here, the question asked is “*Where does E-field strength best predict the stimulation response?*”.

A key requirement for such regression-based TMS mapping is that different cortical locations receive sufficiently distinct E-field patterns across pulses, so that voxel-wise predictors are not excessively correlated. The first implementations achieved this “decorrelation” by quasi-randomly placing the TMS coil above the region of interest, relying on geometric variability in coil pose to generate diverse fields (Numssen et al., 2021; Jing et al., 2023). While this random coil placement procedure worked well for sTMS mappings, selecting random mTMS input currents did not yield successful mappings. Recently, we (Qiu et al., 2026a) and others (Schultheiss & Turi et al., 2025) framed sTMS coil-sequence design as an explicit optimization problem and showed that diversity-aware sTMS coil placement selection rules can substantially improve the efficiency of regression-based mapping, yielding faster convergence of effect maps. While these optimization approaches provide significant improvement in localization performance through their specific coil placement selections, they introduce the need for robotically guided TMS (Kantelhardt et al., 2009; Campos & Rondinoni et al., 2026; Souza et al., 2018; Matsuda & Souza et al., 2025; Matsuda & Souza et al., 2024; Bai et al., 2025) to allow for precise and fast repositioning for a large number of coil placements – a necessity that could potentially vanish with the use of mTMS. To this end, the first aim of this study was to test whether determinant-optimal (D-optimal) current selection would allow for the use of mTMS for voxel-wise TMS mapping.

To do so, we extended our recent framework for determinant-based, kernelized selection of informative E-field patterns in sTMS (Qiu et al., 2026a) from discretized sets of coil placements to the continuous mTMS current space. We used a D-optimal design objective to construct sequences of mTMS current patterns whose induced cortical E-fields are as dissimilar as biophysical constraints permit, and we benchmark these sequences against random current baselines in virtual voxel-wise motor mapping. Specifically, we maximize the determinant of the TMS-induced E-field Gram matrix within a region of interest across multiple pulses, following the D-optimal design literature (Saito et al., 2021). Building on our previous work (Qiu et al., 2026a), we show that this determinant-based optimization of the Gram matrix enables efficient and effective greedy selection of mTMS current vectors, in line with recent fast greedy D-optimal sensor selection algorithms (Joshi & Boyd, 2009).

So far, regression based TMS mapping is based on individual E-field computations. This requires the acquisition of MRI scans for each participant or patient, as well as segmentation and meshing to build individual volume conductor models. The TMS-induced E-field in the head consists of two additive components: the primary field, which is directly proportional to the temporal derivative of the magnetic vector potential *d***A**/*dt*, and the secondary component caused by volume conduction according to Ohm’s law. While the primary field only depends on the geometry of the TMS coil and its position relative to head, the secondary field also depends on the individual head and brain morphology and the associated tissue conductivities. Importantly, for TMS the primary field, which is independent of the individual morphology, dominates the total E-field (Opitz et al., 2011; Stenroos & Koponen, 2019; Wang et al., 2024). Therefore, the second aim of this study was to test whether the gain in accuracy from using E-field models based on individual anatomy (*head-space*) over *d***A**/*dt* models from idealized spherical models without individual anatomy (*coil-space*) outweighs the higher costs of MRI acquisition and individual head modeling.

To this end, we contrast a subject-independent coil-space, based only on the primary field, with a subject-specific head-space, based on the full cortical E-fields from individual head models. This allows us to directly test whether and to what extent we can achieve reasonable mapping accuracy without the need of subject- or patient-specific MRI scans, head models, and field computation.

## 2 Methods

### 2.1 Participants

We used the same *in-silico* cohort as in our previous mTMS focality study (Numssen & Martin et al., 2026). Briefly, individual head models (SimNIBS v4.5, Thielscher et al., 2015) were constructed based on structural MRI data for nine healthy adults (Cam-CAN dataset, https://cam-can.mrc-cbu.cam.ac.uk/dataset/, Shafto et al., 2014), spanning a broad adult age range (mean ± SD age: 53.1 ± 26.80) and screened to be free of neurological or major psychiatric disorders.

### 2.2 TMS coils and arrays

We used three mTMS coil array models representing complementary design principles for electronic E-field steering and two sTMS coil models as a baseline comparison (see Fig. S1):

#### 5-channel planar array (*planar-5*)

A surface array consisting of five planar coils stacked and assembled in a single case, with two figure-of-eight coils, two cloverleaf coils, and one round coil. This design reproduces figure-of-eight–like stimulation patterns, and allows for translating and rotating the E-field peak on the cortical surface within a 30–40 mm diameter space (Fig. 2; Nieminen & Sinisalo et al., 2022).

**Figure 2:**
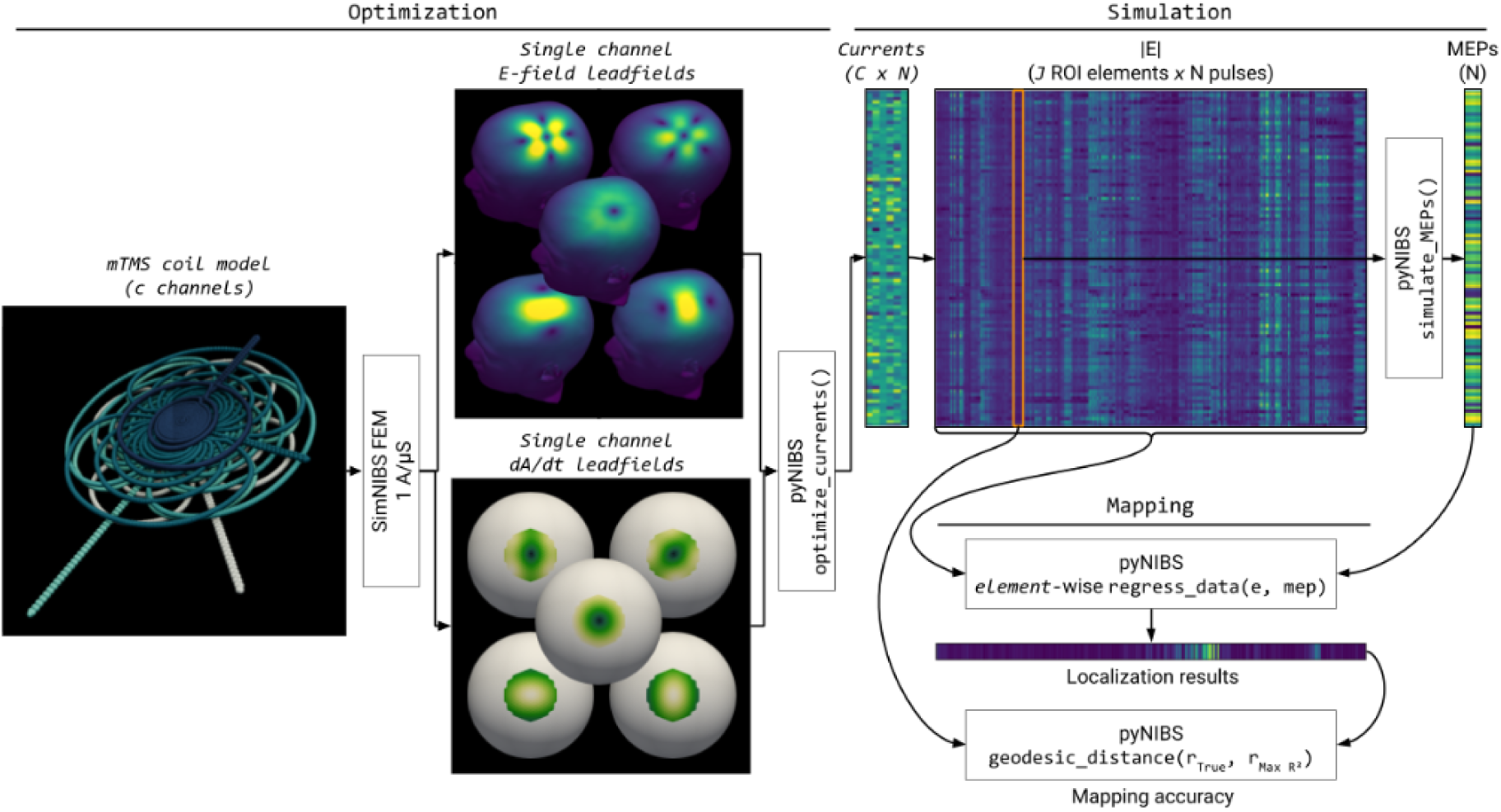
Simulation and mapping pipeline. Starting from an mTMS coil model with *C* channels, we computed single-channel lead fields either as subject-specific cortical ∣**E**∣ patterns (*head-space:* realistic head meshes) or as *coil-space d***A**/*dt* patterns on a spherical surface. To these lead fields we applied D-opt RBF optimization to select stimulation sequences with low pattern redundancy. The resulting current matrices were then mapped through the E-field lead fields to obtain ∣**E**∣ patterns across *J* ROI elements and *N* pulses, which served as input to MEP simulations. Finally, we performed voxel-wise regression ∣**E**∣ and simulated MEP amplitudes to test localisation accuracy.

#### 6-channel spherical array (*spherical-2x3*)

A modular design comprising two three-axis units, each with three circular coils oriented along orthogonal x, y, and z axes. This geometry prioritizes flexible orientation control and modular design (Navarro de Lara et al., 2021).

#### 12-channel spherical array (*spherical-4x3*)

An extension of the spherical 2x3 design with four three-axis units (12 channels total) arranged in a square pattern. The increased channel count allows richer superposition patterns and improved controllability of the E-field distribution at the cost of higher demands on current-rate optimization and hardware constraints (Navarro de Lara et al., 2021).

#### Single channel TMS coils

Additionally, we performed the optimization for two traditional single-channel TMS coil models: a 35 mm and a 65 mm figure-of-eight coil (MagVenture MCF-B35 and MagVenture MCF-B65, respectively; Drakaki et al., 2022).

### 2.3 Subject-specific E-field simulations

#### 2.3.1 Head models

Head models were generated as in Numssen & Martin et al. (2026). T1- and T2-weighted MR images were segmented into multiple tissue classes (gray matter, white matter, cerebrospinal fluid, compact and spongy bone, scalp, eyeballs, and additional tissues where available) using the SimNIBS 4.5 charm pipeline (Puonti et al., 2020) and Freesurfer (Fischl, 2012). High-resolution finite-element meshes were constructed with cortical element volumes on the order of 0.15 mm³. E-fields were simulated in the quasi-static regime using SimNIBS 4.5 and pyNIBS (v0.2026.1; Numssen et al., 2021), assuming unity current rate (1 A/µs) in each channel. For all analyses, we extracted the E-field at the gray-matter mid-layer to reduce boundary artefacts.

#### 2.3.2 Regions of interest

We focused on a left-hemispheric somato-motor region of interest (ROI) encompassing primary motor cortex (M1), primary somatosensory cortex (S1), and dorsal premotor cortex (PMd). The ROI was defined on a standard template (*fsaverage*) and mapped to individual cortical surfaces, then restricted to the gray-matter mid-layer for E-field extraction. This ROI has been used previously for TMS motor mapping (Numssen et al., 2021; Schultheiss & Turi et al., 2025) and mTMS focality optimization and provides a well-characterized testbed for evaluating stimulation patterns.

#### 2.3.3 Coil placements

For each subject and mTMS coil array, we defined a single array placement above the motor ROI. The array center was positioned on the scalp over the hand-knob region in the left precentral gyrus following the placement strategy used in Numssen & Martin et al., 2026. This placement was chosen such that the array covered the entire ROI. No further repositioning was performed within a subject: all stimulation patterns for a given array were generated purely by changing the channel-wise current change rates. This reflects the intended use-case of mTMS as an electronically steerable system. For the sTMS coils, we pre-computed a large set of candidate coil placements (positions and orientations) above the motor cortex ROI (Qiu et al., 2026a), yielding 1488 coil placements per subject and sTMS coil.

#### 2.3.4 Lead-field computation

For each subject and mTMS coil array, we computed channel-wise *E-*lead-fields in the realistic head model ROI (head-space). For a given array with C channels, each channel c was driven with a unit current rate (1 A/µs) (Wang et al., 2024), and the resulting E-field *E_c_*(*r*) was simulated using SimNIBS 4.5 and pyNIBS. We extracted the three-dimensional E-field vector at all elements of the gray-matter mid-layer motor ROI, then concatenated the components into a column vector. Stacking these vectors across channels yielded a lead-field matrix *L* ∈ ℝ^(3*V*)×*C*^, where V is the number of ROI elements and *C* ∈ {5, 6, 12} is the number of channels for the respective array. Under the quasi-static approximation, the E-field induced by an arbitrary current-rate vector I ∈ ℝ*^C^* is given by the linear superposition *E* = *LJ*, which was evaluated reduced to the magnitude ∣**E**∣. Mutual inductance between channels was neglected, consistent with prior modeling of these arrays (Numssen & Martin et al., 2026).

For the sTMS coils, we computed the individual E-field for each of the 1488 candidate coil placements for 1 A/µS and, subsequently, upscaled the E-field to a reasonable stimulation strength, yielding about 100 V/m in the motor cortex ROI.

### 2.4 Coil-space *d***A**/*dt* lead fields

In addition to the head-space, i.e. subject-specific, mTMS E-lead-fields, we constructed purely coil-space representations based on the coils’ *d***A**/*dt* on an idealized sphere, where *A* denotes the magnetic vector potential of the coil array. The term *d***A**/*dt* captures the inductive, divergence-free component of the TMS E-field generated by the time-varying coil currents, i.e., the part of **E** that is present before Ohmic volume conduction in the head redistributes the field via the scalar potential. This coil-space field therefore depends only on coil geometry and drive currents and not on individual head and brain morphology; if optimizing current patterns in this space already yields “good enough” mapping results, then mTMS sequence design could, in principle, be done once per coil array rather than per subject.

We generated a triangulated sphere (radius 75 mm) and defined a circular ROI (radius 25 mm) around the “north pole” to mimic a local cortical patch under the mTMS coil array. Each multichannel coil model was placed above this ROI at a realistic coil–brain distance. For each channel, we computed and exported *d***A**/*dt* for all ROI triangles with SimNIBS, resulting in a channel-wise *d***A**/*dt* tensor over the spherical ROI. We saved the sphere geometry and the channel-wise *d***A**/*dt* values to use them as an alternative design space for our current optimization algorithm. After mTMS current optimization based on *d***A**/*dt*, we used these coil-wise current patterns to generate subject-wise E-fields and MEPs, consistent with the head-space optimization scheme. By comparing current patterns optimized in this spherical *d***A**/*dt* space with those optimized using subject-specific E-field lead fields, we could directly test whether coil-space optimization is sufficient or whether individualized, head-space E-field information is necessary for optimizing mTMS sequences in regression-based mapping.

### 2.5 Optimization of mTMS patterns

#### 2.5.1 Kernelized D-optimal objective

We optimized mTMS current patterns to maximize a determinant-based diversity measure of ROI E-field magnitudes, using the same kernelized D-optimal criterion as in our previous work (Qiu et al., 2026a), adjusted from a discrete version for sTMS coil placements to a continuous-current-based version for mTMS: For a set of *N* stimulation patterns, we assembled the voxel-wise E-field magnitude in the motor ROI into a matrix |*E*| ∈ ℝ*^N^*^×*V*^, with one row per pulse and one column per ROI element. From this matrix, we constructed an RBF kernel

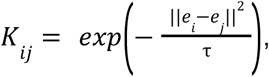

where *e_i_* denotes the i-th row of ∣**E**∣, *τ* is estimated from the median of sampled pairwise squared distances. The design objective is the kernelized D-optimal log-determinant

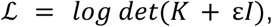

which favors sets of pulses whose ∣**E**∣ patterns span a large “volume” in feature space and are therefore different to one another. Here, *ε* is a small ridge term for numerical stability and ℒ is evaluated via a Cholesky factorization for numerical stability (see Qiu et al., 2026a for details). For the coil-space optimization, the *d***A**/*dt* matrix was used instead of **E**, as described above.

#### 2.5.2 Greedy continuous current optimization

We treated the currents for each pulse as continuous variables and applied a greedy, pulse-by-pulse optimization in current space. Let *L_x_*, *L_y_*, *L_z_* ∈ ℝ*^V×C^* denote the ROI-restricted lead-field components for an array with *C* channels. For a new current vector *I_new_* ∈ ℝ*^C^* and an already selected set of pulses *I_sel_* ∈ ℝ*^C×k^*, we compute:

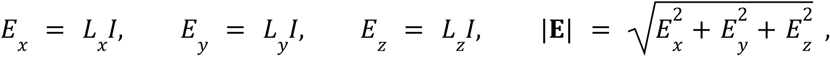

clip |**E**| to within [50 *V*/*m*, 200*V*/*m*] as a proxy for the sigmoidal E-MEP I/O-curve, and form the updated |**E**| matrix over the *k+1* pulses. From this matrix we build the RBF kernel and compute the log-determinant; for each new pulse we minimize the negative log-det value with respect to *I_new_*. Optimization is performed with L-BFGS-B (Zhu et al., 1997) in the continuous current space, subject to simple box constraints |*I_c_*| ≤ *I_max_* for each channel. This yields a greedy, continuous D-optimal design directly on |**E**| in the motor ROI.

#### 2.5.3 Optimization settings

The first pulse is initialized by a separate heuristic that balances high average stimulation strength against current costs. Specifically, we draw random current vectors from a uniform distribution over [− *I_max_*, *I_max_*]*^C^* with I_max_ = 150 A/µS, evaluate the mean ROI |**E**| and a small quadratic current penalty for each, and select *score*(*I*) = *mean*(|**E**(*I*)|) − 10^−4^|*I*|^2^ as the criterion to choose the initial pulse. Subsequent pulses are then optimized via L-BFGS-B from multiple random starts, using the negative kernel log-det objective described above. After each newly accepted pulse, we recompute |**E**| for all selected pulses, re-estimate *τ* from the current |**E**| matrix, rebuild the kernel, and track the resulting log-determinant to monitor decorrelation progress (pynibs.optimization.multichannel.dopt_objective_single_pulse()). To reduce the risk of local minima, each greedy step uses a multi-start strategy, with 20 restarts whenever a better solution is discovered. For each subject, mTMS coil array, and ROI subsampling, we designed sequences of length *N* ∈ {5, 25, 50, 100}, generated matched random-current sequences as baselines for subsequent virtual mapping analyses, and repeated this optimization 20 times with different seeds to yield a large set of optimization results.

### 2.6 Virtual MEPs and mapping readout

To create regression-based virtual motor-mapping data, we simulated MEPs originating from multiple cortical targets using the dual-variability-source (DVS) model of Goetz et al. (2014), implemented in pyNIBS (get_mep_virtual_subject_DVS()) as a realistic IO-curve generator. For each subject and optimization condition, we randomly selected 50 ROI elements as ground-truth motor targets, sampling across gyral crowns, rims, and sulcal walls to reflect typical motor-mapping scenarios. For each optimized and random current sequence (E-field-based: sTMS and mTMS; *d***A**/*dt*-based: mTMS only), we extracted the local E-field magnitude at the target element for every pulse and passed this to the dual-variability-source model to generate realistic MEPs.

We simulated three levels of response variability, parameterized as noise levels σ ∈ {0, 0. 1, 0. 2} applied to σ_x_ and σ_Y_ of the DVS model, covering deterministic to strongly noisy regimes, including and surpassing realistic noise levels seen in motor cortex studies (e.g. Numssen et al., 2021). For each combination of target (50), noise level (3), and optimization space (E-field vs *d***A**/*dt*), we generated 20 independent realizations (random seeds) per subject (9) and mTMS coil array (3) yielding 50 × 3 × 2 × 20 × 9 × 3 = 162, 000 mTMS MEP series, each for 5, 25, 50, 100 stimulations. As a baseline comparison, we performed a random-current selection approach (162,000 MEP series), parallelizing our sTMS regression approach (Numssen et al., 2021), where the sTMS coil was positioned randomly above ROI for each pulse. For sTMS comparison, the same optimized and random current selection procedure was performed for the E-field optimization space only. This factorial design allowed us to separate effects of sequence design from pure response noise.

For the final readout, we used two mapping approaches: A linear-fit-based and a mutual-information-based hotspot identification. For each virtual experiment, we assembled the E-field magnitude (and *d***A**/*dt* magnitude, respectively) across all ROI elements and stimulations into a matrix |**E**| ∈ ℝ*^N^*^×*V*^ (and|*d**A***/*dt*| ∈ ℝ*^N^*^×*V*^) and treated the simulated MEP amplitudes as the target variable. Due to the inherent smoothness of the E-field, we subsampled the E-field ROI to 4,000 equi-distant elements to speed up mapping performance. We then computed a linear regression and the mutual information between the MEP vector and the **E** and *d***A**/*dt* pulses at each ROI element, applied to |**E**|. The ROI element with maximum R² score or maximum mutual information, respectively, was taken as the estimated hotspot, and geodesic distances between true and estimated hotspots were collected across targets, noise levels, and sequence designs, to quantify how well different mTMS optimization strategies support regression-based recovery of the true stimulation site.

In summary, we optimized mTMS current patterns in two design spaces, a coil-space based on *d***A**/*dt* and a head-space based on cortical |**E**|, for different sequence lengths (5, 25, 50, 100 pulses) across sTMS coils and mTMS coil arrays. For each optimized sequence, we then propagated the selected currents through the subject-specific E-field lead fields, simulated MEPs from 50 randomly chosen ROI locations, and evaluated how well linear- and mutual-information-based voxel-wise mapping recovered the true hotspots.

## 3 Results

### 3.1 sTMS vs mTMS optimization

D-optimal coil placement selection for sTMS and current optimization for mTMS both substantially reduced E-field correlation and improved hotspot localization, confirming the successful adaptation of determinant-based sequence design to the multichannel current space (Fig. 3 shows an exemplary subject at 25 pulses). Across all coil types and TMS systems, increasing MEP variability from noise-free responses (noise level 0) to more realistic, noisy regimes (noise level 0.2) degraded mapping accuracy for both linear regression and mutual-information readouts, so that longer stimulation sequences up to 100 pulses were required to reach stable localization performance (linear fit: Fig. 4, rows; mutual information: Fig. S2).

**Figure 3:**
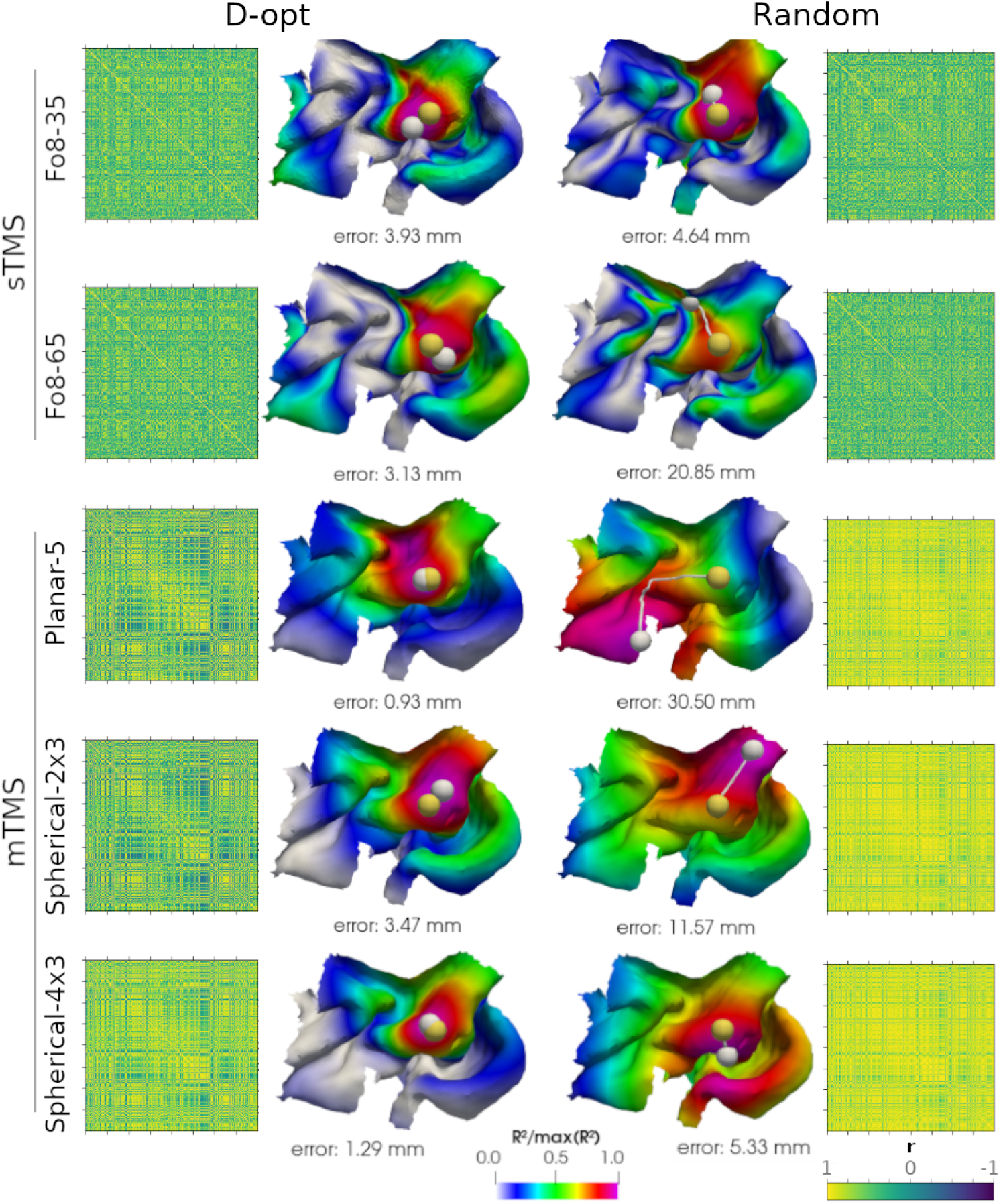
Optimizing E-fields improves localization accuracies for sTMS and mTMS. Left: D-optimal coil placement selection (sTMS; top two rows) and current optimization (mTMS; bottom three rows). Right: Random coil placements and current selections. Random current selection (mTMS) yields high autocorrelation and poor mapping accuracy, whereas determinant-based optimization yields high accuracy for sTMS and mTMS. Exemplary subject with true hotspot at gyral crown (yellow sphere), 25 pulses, linear fit, noise level 0.1. White sphere: identified location. Correlation matrices: E-field correlation in the motor cortex ROI across 25 pulses (4,000 ROI elements).

**Figure 4:**
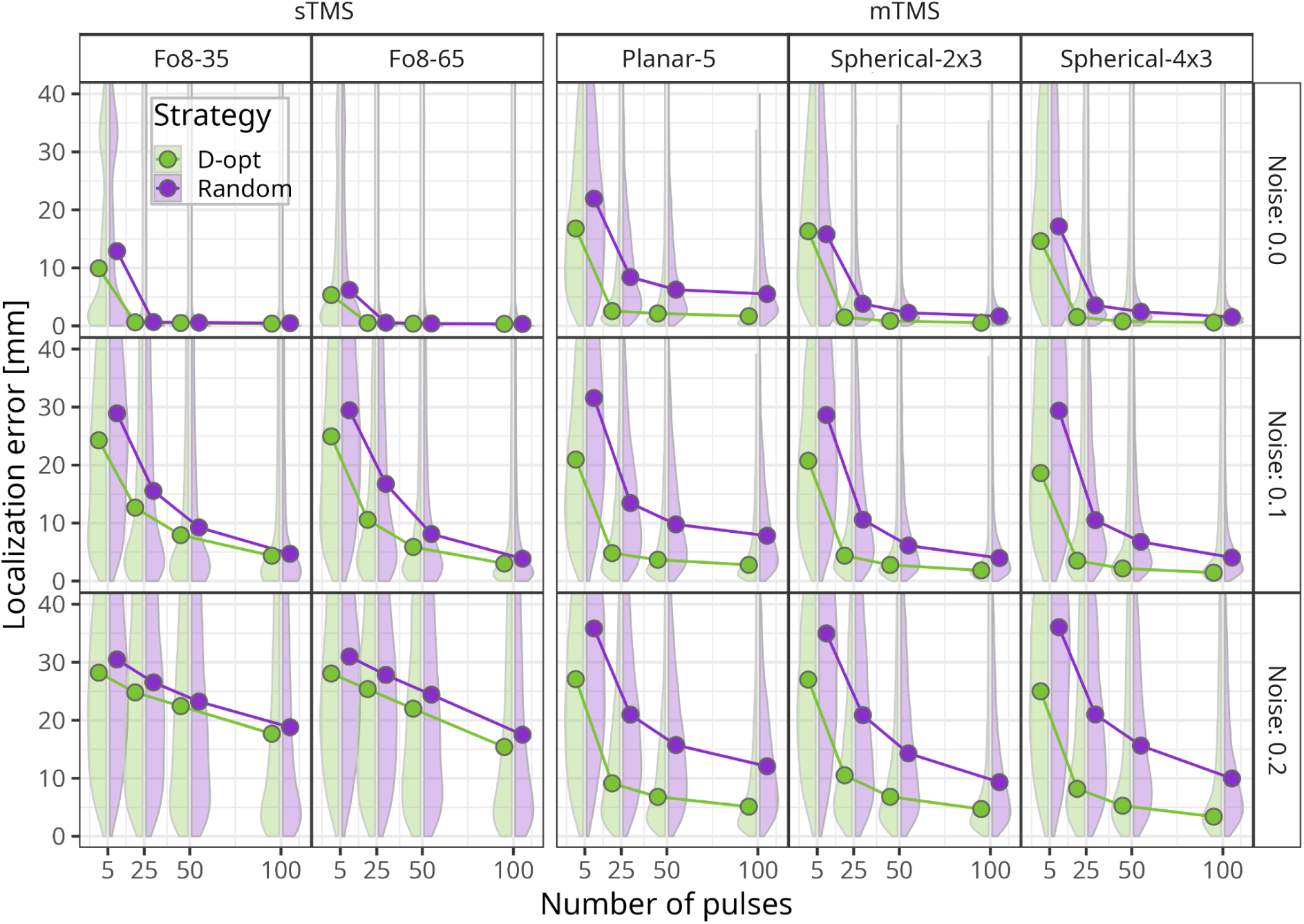
E-field-aware optimization: linear fit mapping accuracy across sTMS and mTMS coil types and MEP noise levels. For each of five coil configurations (columns; two figure-of-eight sTMS coils (35 mm and 65 mm diameter); three mTMS coils: planar-5, spherical-2×3, spherical-4×3), the mean geodesic distance between the true and detected target location (y-axis) is plotted against the number of TMS pulses (5, 25, 50, 100; x-axis). Rows: levels of MEP noise (0.0, 0.1, 0.2). Green lines and markers: D-optimal (D-opt) coil placement (sTMS) or current optimization (mTMS) strategy and violet for random selection. Violet: random coil placement/current selection. Simulations were performed for 9 individual motor cortex ROIs, with 50 random targets localized, fitted with a linear function.

Under noise-free conditions, random and optimized sTMS coil placements yielded near-perfect localization after as few as 25 pulses, whereas random mTMS currents clearly underperformed their optimized counterparts. This larger optimization gain for mTMS persisted with increasing noise: at medium MEP noise (0.1), optimized mTMS sequences already produced successful mapping across the ROI with 25 pulses (linear-fit localization error (geodesic distance), median ± SD: planar-5: 4.91 ± 6.73 mm; spherical-2×3: 4.61 ± 7.43 mm; spherical-4×3: 3.63 ± 6.41 mm), in contrast to substantially higher errors for random currents (planar-5: 14.76 ± 14.85 mm; spherical-2×3: 12.31 ± 13.1 mm; spherical-4×3: 12.20 ± 13.30 mm). For the same number of pulses, mTMS optimization also outperformed sTMS coil placement optimization (25-pulse linear-fit errors: Fo8-35: 16.47 ± 20.21 mm; Fo8-65: 14.59 ± 20.01 mm), with a robust statistical advantage for mTMS over sTMS (F(1, 90520) = 24742.92, *p* < .001). In the high-noise regime (0.2), this pattern was further amplified: even after 100 pulses, sTMS sequences did not support reliable localization (Fo8-35: 22.37 ± 20.84 mm; Fo8-65: 20.71 ± 20.65 mm), whereas all three mTMS arrays achieved high accuracy, with median errors below 6 mm (planar-5: 5.74 ± 7.18 mm; spherical-2×3: 5.80 ± 8.60 mm; spherical-4×3: 4.16 ± 7.28 mm).

Overall, linear regression (Fig. 4) and mutual information (Fig. S2) produced comparable error profiles, with linear fits slightly favoring sTMS and mutual information slightly favoring mTMS (Fig. 5; Figs. S4 & S5). Taken together, these results demonstrate that mTMS current optimization enables robust, high-resolution localization of virtual MEP sources in motor cortex ROIs, matching sTMS performance in low-noise conditions and clearly surpassing it when signal-to-noise ratios are reduced, despite relying on a single fixed array placement rather than pulse-wise coil repositioning.

**Figure 5:**
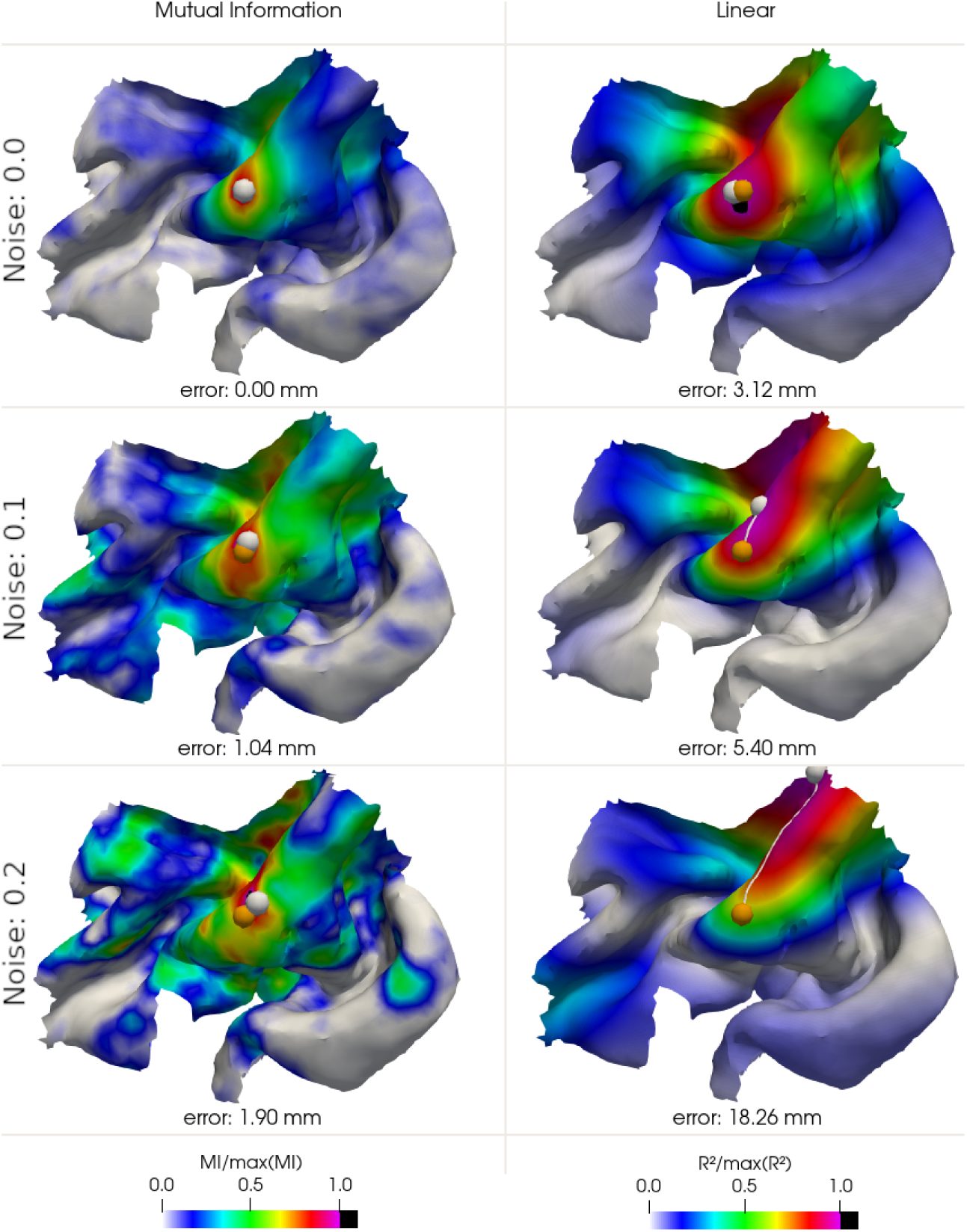
Effect of noise on recovery accuracy across mutual information and linear regression. Exemplary mapping results after 50 D-optimal pulses with the planar-5 mTMS system for a gyral crown target in the center of the ROI (yellow sphere). White sphere: identified location. Columns: Mutual information regression (left) and linear regression (right). Rows: Different MEP noise levels.

### 3.2 Coil-space versus head-space optimization

Head-space optimization of mTMS input currents directly targets the quantity of interest: the induced cortical E-field within the ROI. While conceptually straightforward, this strategy requires subject-specific current optimization, imposing demands on time, infrastructure, and computational resources. We therefore asked whether a purely coil-space scheme, optimizing currents based on *d***A**/*dt* as a proxy for the primary E-field, could serve as a more resource-efficient alternative. Because this coil-space optimization depends only on array geometry, it can be performed once per coil model rather than per participant. Across mTMS systems and noise levels, coil-space current optimization (orange; Fig. 6) consistently outperformed random current selection (violet; Fig. 6) (*F(1, 1989258)* = 34417.29, *p* < .001) but led to significantly worse mapping accuracy than head-space current optimization (*F(1, 1814508)* = 4692.23, *p* < .001), with larger effect sizes of coil-space > *random* (Eta² = 0.02) than head-space > coil-space (Eta² = 2.58e-03). In medium (0.1) and high (0.2) noise regimes, head-space (green) and coil-space (orange) designs yielded similar localization accuracies, with a small advantage of head-space optimization for sequences up to 50 pulses. For 100 pulses, we did not observe any detectable difference in mapping performance between the two approaches, indicating that coil-space designs can indeed capture most of the available sequence-optimization benefits when enough mTMS pulses are used.

**Figure 6:**
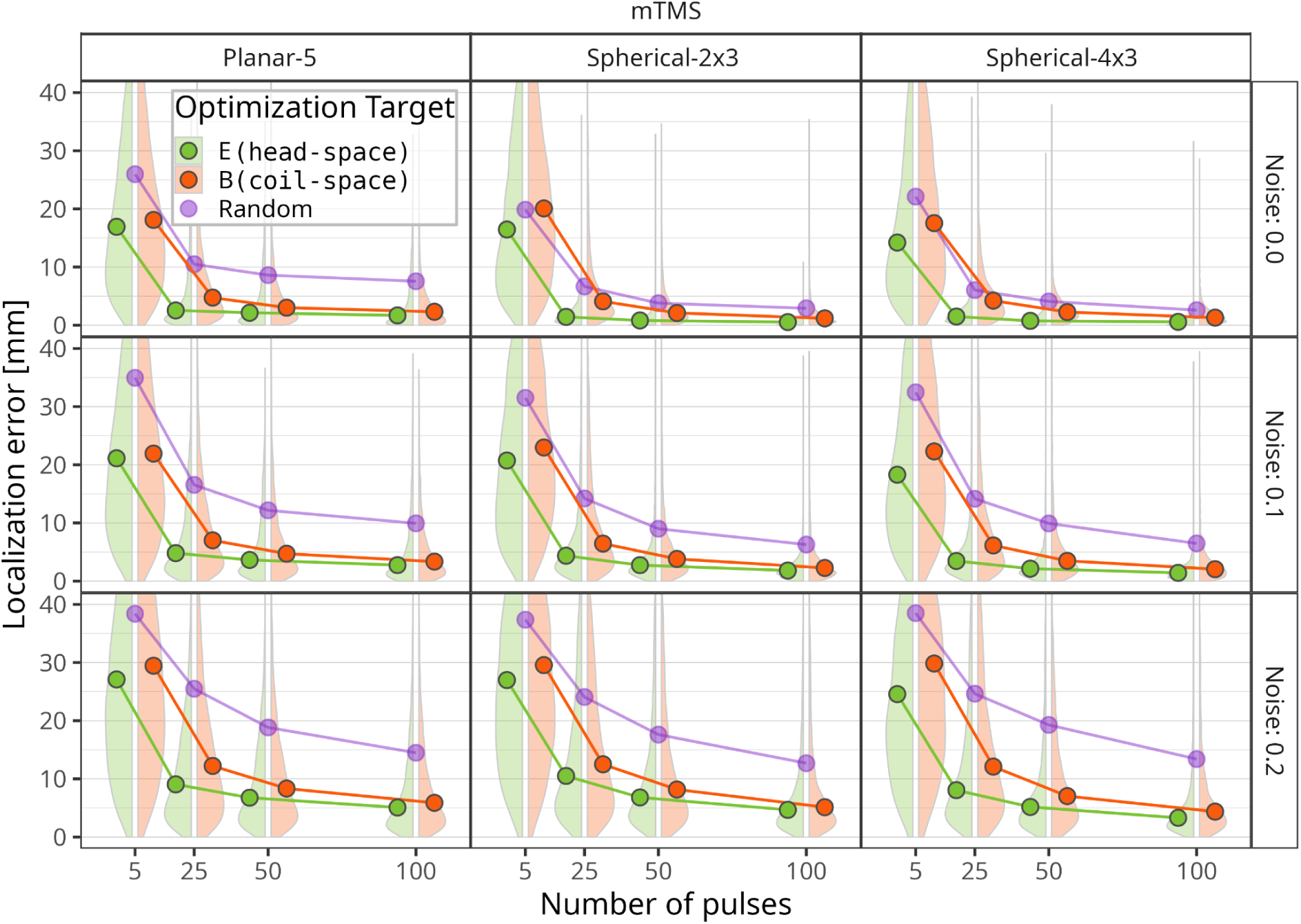
Leveraging individual stimulation patterns improves mapping. Current-pattern optimization was performed either based on the individual E-field patterns (green, ‘*head-space’*) or on the magnetic field of the coil (orange, ‘*coil-space*’), which does not take individual factors such as gyrification into account. While individual-specific optimization yielded the best mapping results, coil-space optimization yielded consistently better mapping performance than random currents.

### 3.3 Effects of cortical source location

Cortical source location systematically affects voxel-wise mapping accuracy, with the largest gains from optimized current sequences emerging for deeper motor targets. For mTMS, random current selection yielded relatively poor and depth-dependent localization across all skin–cortex distances (Fig. 7, bottom row). Consistent with previous work on sTMS sequence design (Qiu et al., 2026a), superficial sources with skin-cortex distances up to approximately 20 mm could be recovered reliably from random coil placements, with particularly good performance for the smaller figure-of-eight coil (Fo8-35). D-optimal sTMS coil placement further improved accuracy for the larger Fo8-65 coil, but even optimized sTMS sequences failed to achieve robust localization for targets deeper than about 25 mm. For these deeper sources, D-optimal mTMS currents produced strikingly low localization errors across the entire motor ROI, including targets with skin–cortex distances up to roughly 35 mm, and performance improved with increasing channel count in the spherical arrays (2×3 and 4×3), consistent with richer spatial variability afforded by more diverse coil geometries.

**Figure 7:**
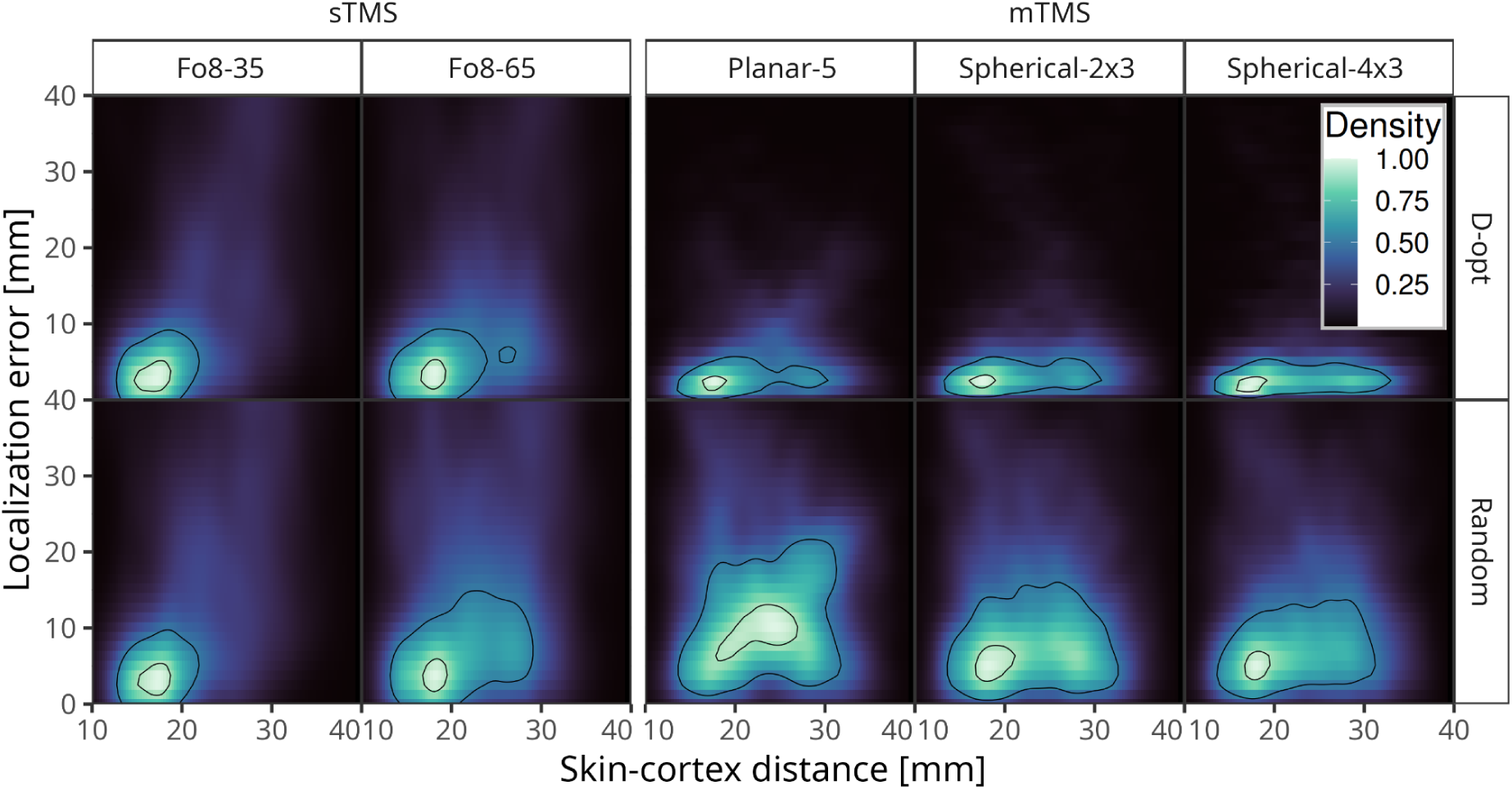
Optimal E-field selection improves localization of deep targets. Both for sTMS (left columns) and mTMS (right columns) the D-optimal E-field selection (top row) yields better performance than the random approach (bottom row). Especially for deeper targets (skin-cortex distance above ∼25 mm) the localization error is lower for the optimal solution. Linear fit. Black contour lines: 50 and 90 percentiles.

Besides depth, we asked how mapping accuracy changed for cortical targets with varying distance to the ROI center (Fig. S3). Because sTMS coils were repositioned for each pulse while mTMS arrays were kept fixed over the ROI center, effects of target distance from the ROI center potentially affect mapping accuracy differently across mTMS and sTMS systems. As expected, sTMS mapping accuracy was essentially independent of ROI-center distance for both random and optimized coil placements, confirming that the sTMS candidate-placement grid adequately covers the ROI. Strikingly, mTMS showed uniform, superior performance across ROI-center distances, indicating accurate localization throughout the full motor ROI despite a single, static array placement; again, D-optimal mTMS current design provided larger accuracy gains than D-optimal sTMS coil placement compared to random mappings.

Together, these results demonstrate that sequence optimization in mTMS primarily mitigates the depth-related limitations of standard TMS while preserving high localization fidelity across the lateral extent of the motor cortex.

## Discussion

Here, we adapted determinant-based, kernelized sequence optimization from single-coil placements (Qiu et al., 2026a) to the continuous current space of multichannel TMS (mTMS), to test whether electronically decorrelating mTMS-induced E-fields improves voxel-wise, regression-based recovery of virtual MEP sources. Across three mTMS geometries and a wide range of pulse counts, response noise levels, and cortical depths, D-optimal current design consistently and substantially outperformed random current selection, and, critically, outperformed D-optimal *coil-placement* selection for standard, single-coil TMS (sTMS) at matched pulse numbers. This advantage was modest under artificial, noise-free conditions, where already randomly placed sTMS pulses localized superficial sources almost perfectly within 25 pulses. This advantage increased sharply as response noise increased: at high noise, optimized sTMS sequences failed to converge even after 100 pulses, while all three mTMS arrays reached median errors at or below 6 mm, comparable to a recent Bayesian mTMS hotspot mapping approach (Granö et al., 2025). A similar pattern held for cortical depth: mTMS uniquely recovered targets down to ∼35 mm skin–cortex distance, well beyond the ∼25 mm ceiling of even optimized sTMS.

### mTMS provides more robust localization at higher MEP variability and increased cortical depth

D-optimal design theory predicts that the benefit of maximally decorrelated predictors over unstructured sampling scales with the noise-to-signal ratio of the outcome variable (Joshi & Boyd, 2009): under low noise, almost any sufficiently diverse set of E-field patterns identifies the correct cortical location, but as signal-to-noise reduces, the estimator’s variance becomes increasingly sensitive to the conditioning of the design matrix. sTMS regression mapping is constrained by a single, fixed coil geometry (see Numssen & van der Burght et al. (2023) for a discussion of sTMS focality limitations), which caps how decorrelated its candidate E-field patterns can become. The channel-wise current superposition for mTMS instead allows for the same physical array placement to produce highly distinct ROI-wide field patterns. This is consistent with our finding that the mTMS advantage over sTMS was essentially flat across distance from the array center despite a single, static placement while sTMS accuracy depended only on the coverage of its (1,488-placement) candidate grid.

### Coil-space optimization improves mapping over random current selection

A central and practical question is whether these gains for structure-function mappings require subject- and session-specific optimization. Our coil-space, spherical *d***A**/*dt* optimization, which can solely be performed on the mTMS coil array geometry and, thus, can be precomputed once per coil model rather than once per participant, nearly matched subject-specific head-space performance at 100 pulses, with a significant, but small edge for head-space designs for fewer pulses across all noise levels. Critically, the coil-space optimization consistently outperformed random current selection (across all noise levels and pulse numbers). This mirrors the pragmatic framing of our focality study (Numssen & Martin et al., 2026), and suggests that off-the-shelf, precomputed D-optimal sequence libraries per mTMS array could be distributed without per-subject MRI or FEM overhead, lowering the barrier for adoption while leaving individualized optimization as a resource worth investing in specifically for short, time- or safety-constrained protocols.

### Beyond motor mapping: implications for studies on higher cognitive functions

Voxel-wise TMS mapping was built and validated almost entirely on the MEP as an outcome measure (Opitz et al., 2013; Aonuma et al., 2018; Laakso et al., 2018), specifically chosen as a high-SNR, single-origin readout with a well-characterized monotonic input-output relationship (Weise & Numssen et al., 2020; Numssen et al., 2021; Weise & Numssen et al., 2023; Vetter et al., 2025; Laine et al., 2026). The first attempt to transfer this framework to a genuinely cognitive domain, Jing et al.’s (2023) sTMS mapping of spatial attention in the right IPL, did not find a robust E-field-behavior relationship at the group level, which was attributed to “the challenges of transferring the localization method to cognitive functions with high neural response variability and complex network interactions.” Our simulations offer a concrete, mechanistic reading of that result: reaction-time and accuracy-based behavioral outcomes are intrinsically noisier, single-trial measures than an EMG-recorded MEP, plausibly placing cognitive mapping experiments in or beyond the “noise level 0.2” regime we modeled here, precisely the regime in which even optimized sTMS sequences failed to recover the true site, while D-optimal mTMS remained accurate. If this correspondence holds empirically, the results in Jing et al. (2023) may reflect a statistical ceiling of sTMS-based sequence diversity rather than an absence of a genuine structure-function relationship in attention networks, and mTMS-based voxel-wise mapping could be the missing methodological piece needed to extend motor-cortex mapping development toward attention, language, and executive-function paradigms, domains where hundreds of trials are already routinely collected (e.g., Posner-task designs), matching the “many-pulses, high-noise” regime where our results show mTMS to be most advantageous. This logic is complemented by two recent studies that used single-coil E-field regression to recover physiologically meaningful, comparatively low-SNR neuro-cardiac coupling from prefrontal TMS (Feng & Martin et al., 2026; Feng et al., 2026), showing that regression-based mapping recovers relevant information outside of the motor cortex with enough pulses and clear hypotheses. Combining that generalizability with mTMS’s noise robustness is, in our view, a natural next target.

### A structure-independent mapping tool

A second implication of the depth and ROI-coverage results is of practical relevance: because mTMS achieves this accuracy from a single, potentially manually positioned array rather than hundreds of discrete, precisely tracked coil placements, it removes the dependency on the robotic positioning infrastructure that recent sTMS sequence-optimization work increasingly requires to realize its gains (Qiu et al., 2026a; Matsuda & Souza et al., 2024; Matsuda & Souza et al., 2025; Schultheiss & Turi et al., 2025; Bai et al., 2025). Combined with its uniform performance across both depth and lateral ROI position, this argues for mTMS voxel-wise mapping as a genuinely structure-independent tool: well suited to ROIs that mix superficial and deep, or spatially distributed, targets, which is closer to the reality of most cognitive networks than the largely superficial, well-circumscribed hand-knob target of classical motor mapping. This perspective adopted here also accounts for the slightly larger sTMS localization errors compared to earlier results: previously, we either asked how well the mapping procedure could identify predefined (and mostly superficial) MEP origins (Numssen et al., 2021) or single, specific targets (Qiu et al., 2026a), which allowed us to also optimize sampling of the full input-output curve. Here, in contrast, we asked how well any target within the cortical region of interest could be recovered, by applying exactly one set of TMS pulses to map potentially more spread functions like multi-muscle interactions during directed movements (Tardelli & Souza et al., 2022; Morishita et al., 2025; Gordon et al., 2023).

### Toward a coherent mTMS toolkit and closed-loop mapping

Together with our focality-first framework (Numssen & Martin et al., 2026), this study delineates two complementary poles of mTMS use: focality optimization for confirmatory stimulation of an already-known target, and D-optimal decorrelation for exploratory mapping of unknown structure-function relationships. Because mTMS currents can be reprogrammed electronically within milliseconds, sequence design need not remain fixed in advance; our results suggest that closed-loop schemes, which natively rely on greedy, sequential designs that update the decorrelation objective online as data accumulate could further reduce the number of pulses required, in the spirit of EEG-feedback-guided TMS optimization (Tervo & Nieminen et al., 2022; Qiu et al., 2026b), particularly by allocating additional pulses to cortical subregions where hotspot uncertainty remains high. This would move mTMS mapping from a purely pre-experimental design problem toward an adaptive, sequential one, and dovetails with ongoing clinical mTMS hardware development (Sinisalo & Kahilakoski et al., 2025).

### Limitations

Several simplifications should be resolved before that translation: (i) a single generative noise model (Goetz et al., 2014) cannot capture all physiological noise sources (state-dependent excitability, background EMG, coil-to-cortex drift); (ii) we tested a single, fixed array placement per subject and did not combine electronic current steering with array repositioning, which could further improve accuracy; (iii) we used E-field magnitude only, ignoring the directional sensitivity of the cortical activation function affecting, potentially underestimating response variance especially for low E-field regimes (Jing et al., 2026; Weise & Worbs et al., 2023; Souza et al., 2022; Aberra et al., 2020). Similarly, effects of different pulse waveforms and pulse durations, e.g. biphasic vs monophasic, on neuronal recruitment were ignored; (iv) mutual inductance between mTMS channels was neglected; (v) results are confined to a left-hemispheric motor cortex ROI in nine adults, and generalization to other cortical regions, folding patterns, or clinical populations with altered anatomy remains untested (Gomez-Tames et al., 2018); (vi) the current optimization utilized here was constrained by hardware thermal limits.

### Conclusion

This study is one step in a longer methodological arc: from the original E-field regression approach that first localized cortical muscle representations from deliberately placed sTMS pulses (Weise & Numssen et al., 2020), through efficient nonlinear mapping from random coil placements (Numssen et al., 2021) and a validated, protocolized pipeline (Weise & Numssen et al., 2023), to the first attempt to push this structure-function framework into the cognitive domain (Jing et al., 2023) and toward physiological network mapping (Feng & Martin et al., 2026; Feng et al., 2026). Each step exposed the same bottleneck: the diversity of sTMS E-field patterns is fundamentally limited (Qiu et al., 2026a).

Here we show in silico that electronically steered mTMS currents can be optimized to decorrelate cortical E-fields far more effectively than any achievable coil-placement schedule and that this advantage is largest exactly where mapping is hardest, i.e. in high-noise and deep-target regimes. Additionally, optimizing input currents per mTMS coil model instead per individual head and brain morphology yields sufficient mapping results for many signal-to-noise regimes. Together with our focality-first mTMS framework (Numssen & Martin et al., 2026), this work provides the methodological argument that optimizing mTMS sequences is a promising technique to scale voxel-wise TMS mapping from a motor-cortex proof of concept to a general-purpose tool for mapping the noisier, deeper, and distributed structure–function relationships that define higher cognitive functions.

## Acknowledgements

ON is supported by Scads.AI. ON and KW were supported by the Federal Ministry of Education Germany (Bundesministerium für Forschung, Technologie und Raumfahrt, BMFTR, Grant no. 01GQ2201 to TRK). RQ is supported by the BMFTR (Grant no. 13GW0738A to TK and KW). RM is supported by the Jenny and Antti Wihuri Foundation (Grant No. 00250241). We thank Risto Ilmoniemi for providing information to model the 5-channel planar mTMS coil array and Mohammad Daneshzand, Evgenii Kim, and Aapo Nummenmaa for information on the 6-channel and 12-channel spherical mTMS models.

## Data availability statement

All scripts to generate the FEM and dA/dt data, to perform the analyses are published here: https://gitlab.gwdg.de/tms-localization/papers/mtms_voxmap. Optimization functions are made available via pyNIBS: https://pypi.org/project/pyNIBS/.

