## Supplementary material for "One placement, many fields: robust voxel-wise mapping with multichannel TMS": SI

### mTMS coil arrays

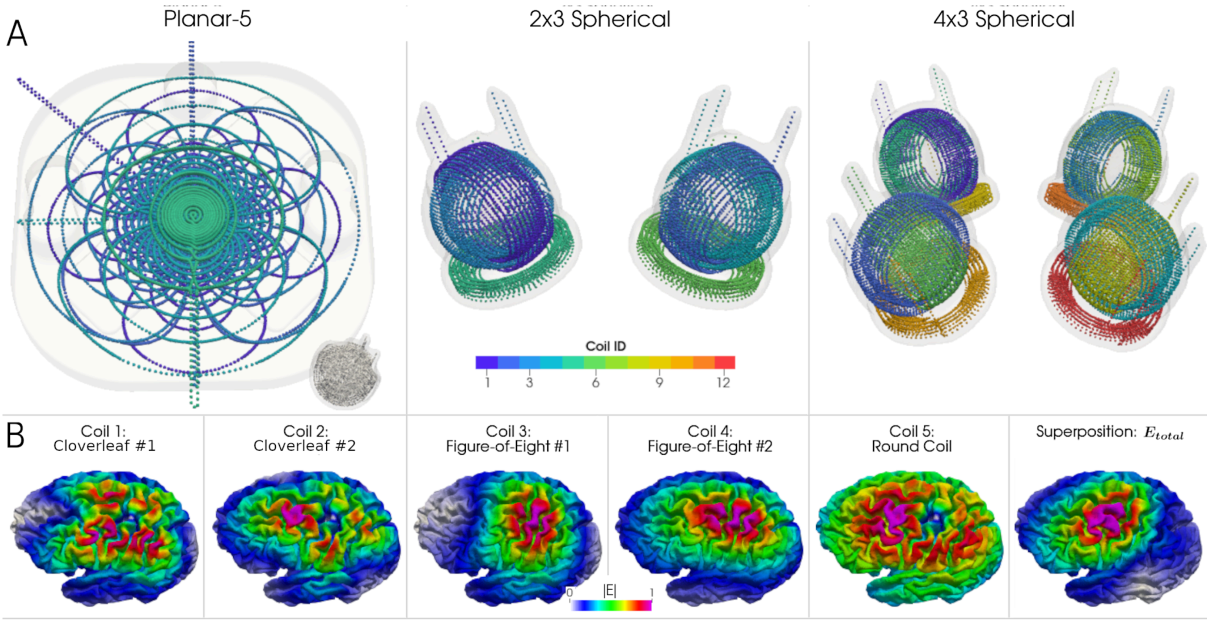


Figure S1: Multichannel TMS (mTMS) coil arrays. A) Coil models for three mTMS coil arrays were constructed: A planar design consisting of five planar coils (left); and two modular, spherical designs, consisting of two (middle) and four (right) three-channel units, respectively, with one circular coil per x-, y-, and z-axis. Inset in left panel: A three-axis unit for size comparison. B) Single-channel E-field examples for the 5-channel planar coil. After multiplying these lead fields with the actual current rates j, i.e., the stimulator output, their vector-superposition yields the total induced E-field $E_{total}$ (right). Figure reproduced from Numssen & Martin et al., 2026.

### Results

#### sTMS vs mTMS

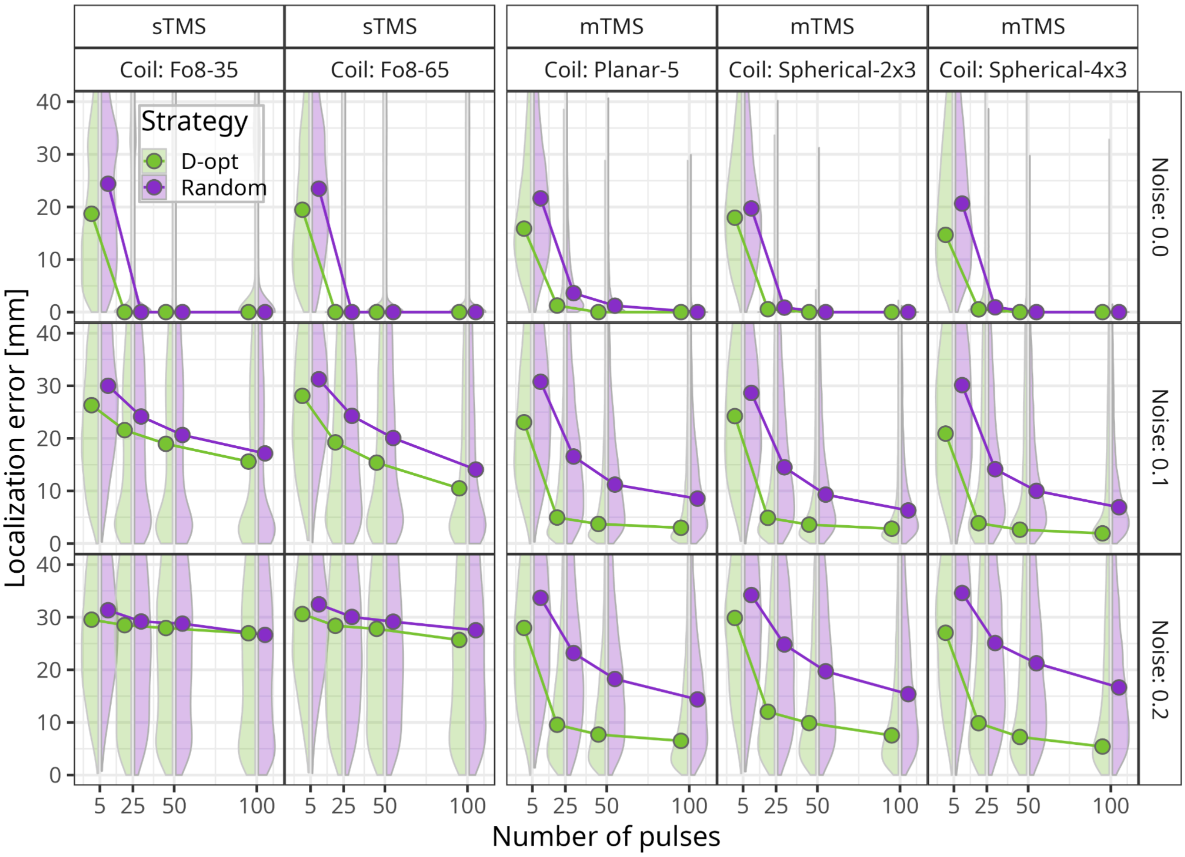


Figure S2: Mutual information. E-field-aware optimization: mapping accuracy across sTMS and mTMS coil types and MEP noise levels. For each of five coil configurations (columns; two figure-of-eight sTMS coils (35 mm and 65 mm diameter); three mTMS coils: planar-5, spherical-2×3, spherical-4×3), the mean geodesic distance between the true and detected target location (y-axis) is plotted against the number of TMS pulses (5, 25, 50, 100; x-axis). Rows: levels of MEP noise (0.0, 0.1, 0.2). Green lines and markers: D-optimal (D-opt) coil placement (sTMS) or current optimization (mTMS) strategy and violet for random selection. Violet: random coil placement/current selection. Simulations were performed for 9 individual motor cortex ROIs, with 50 random targets localized, fitted with mutual information.

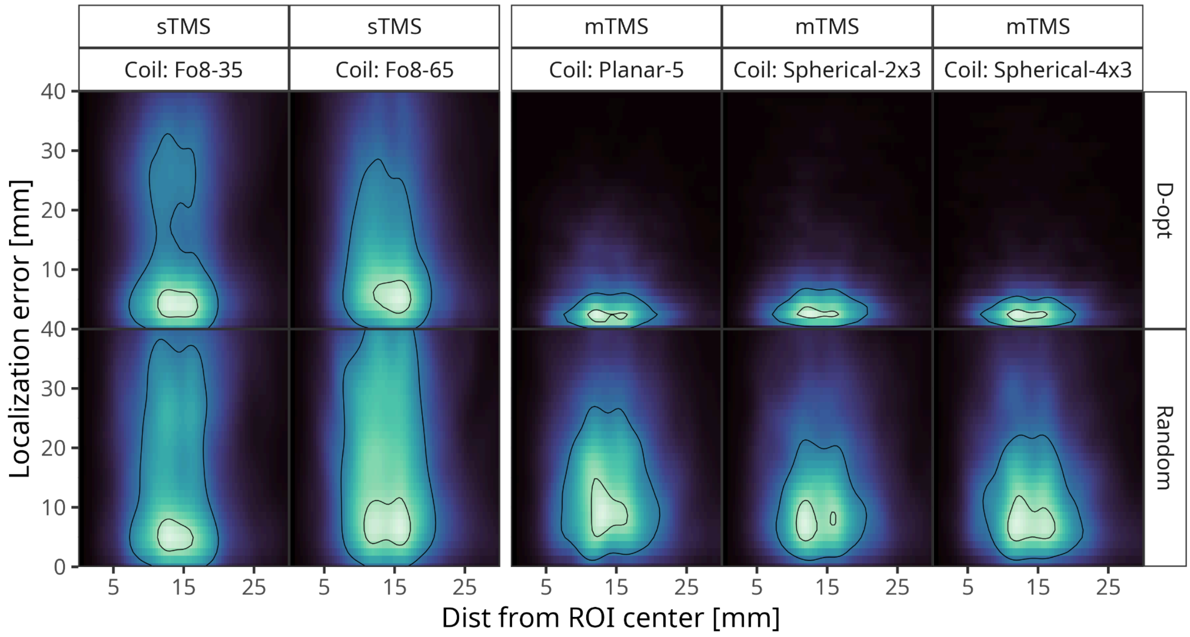


Figure S3. Effects of cortical source location on voxel-wise motor mapping accuracy across sTMS and mTMS coils. Localization error (y-axis; geodesic distance between true and estimated hotspot) is plotted as a function of the target's distance from the ROI center (x-axis) for two single-coil sTMS configurations (Fo8-35, Fo8-65; left columns) and three mTMS arrays (planar-5, spherical-2×3, spherical-4×3; right columns). The top row shows results for D-optimal E-field sequence design, and the bottom row for random current or coil-placement selection. Color maps indicate the density of localization errors across 50 motor targets, 9 subjects, and 25 MEP realizations per condition; white contour lines mark the 50th and 90th percentiles of the error distribution.

#### Mutual Information versus Linear

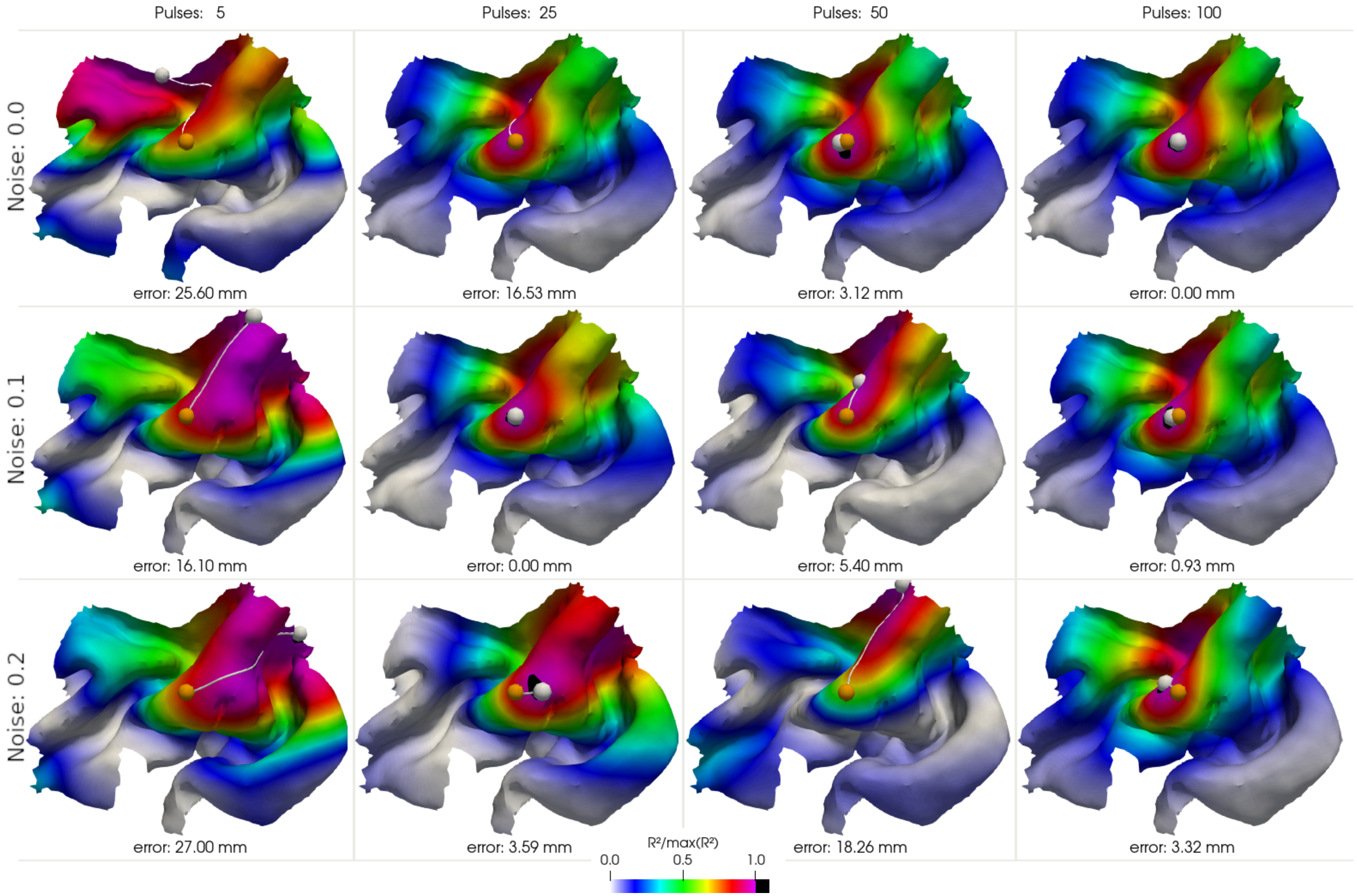


Fig. S4: Linear regression - Effect of noise on recovery accuracy across noise levels and number of pulses. Exemplary mapping results after 5, 25, 50, 100 D-optimal pulses (columns) with the planar-5 mTMS system for a gyral crown target in the center of the ROI (orange sphere). White sphere: detected element (spatial average in case multiple elements share the maximum $R^2$ score (black region). Rows: Different MEP noise levels.

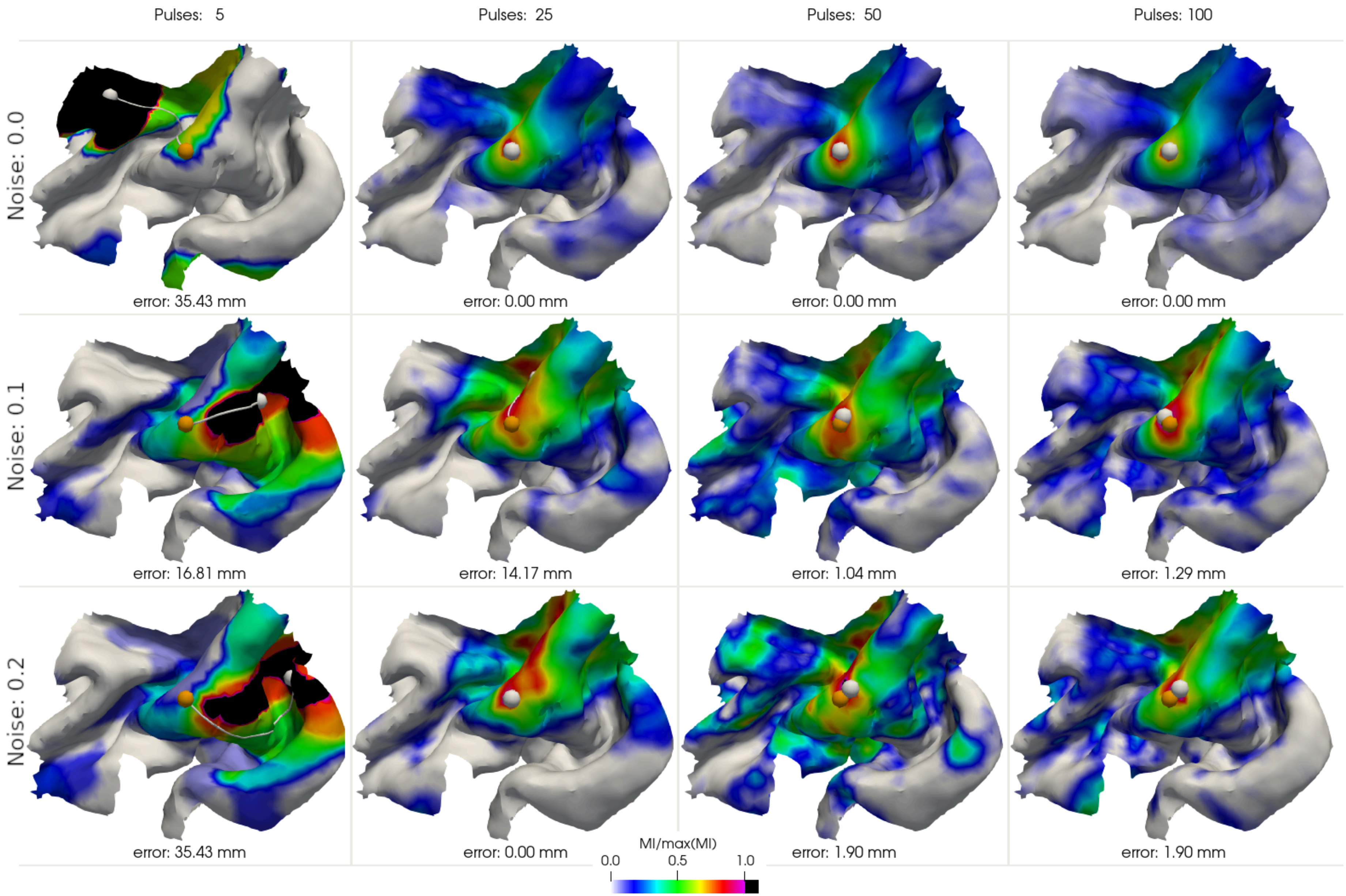


Fig. S5: Mutual information regression - Effect of noise on recovery accuracy across noise levels and number of pulses. Exemplary mapping results after 5, 25, 50, 100 D-optimal pulses (columns) with the planar-5 mTMS system for a gyral crown target in the center of the ROI (orange sphere). White sphere: detected element (spatial average in case multiple elements share the maximum MI score (black region). Rows: Different MEP noise levels.
